# A liquid handling platform for standardised quantification of cell-free enzymatic activity encoded by antimicrobial resistance genes

**DOI:** 10.64898/2026.04.23.720151

**Authors:** Molly Bergum, Bethany Martin, J. Mark Sutton, Simon J. Moore

## Abstract

Antimicrobial resistance (AMR) is a growing global threat to human health, and rapid methods for characterising emerging antimicrobial resistance genes (ARGs) are needed. Here, we develop a semi-automated workflow using cell-free gene expression (CFE) systems to measure the activity of two ARGs encoded on plasmid DNA that produce rifampicin-inactivating and gentamicin-inactivating enzymes. We validated the use of a small benchtop Myra liquid handling system compared to manual pipetting, with no statistical differences observed. After optimising the pre-incubation time of ARGs and dispensing protocol, expression of *aac(3)-IIa* increased the half-maximal inhibition concentration (IC_50_) of gentamicin by over 150-fold, while *arr-3* increased the IC_50_ of rifampicin by approximately 20-fold compared to controls. Future work could extend this platform to characterise novel ARGs identified through genomic surveillance or rapidly profile activity of new or derivative antibiotics.

**Graphical Abstract:** 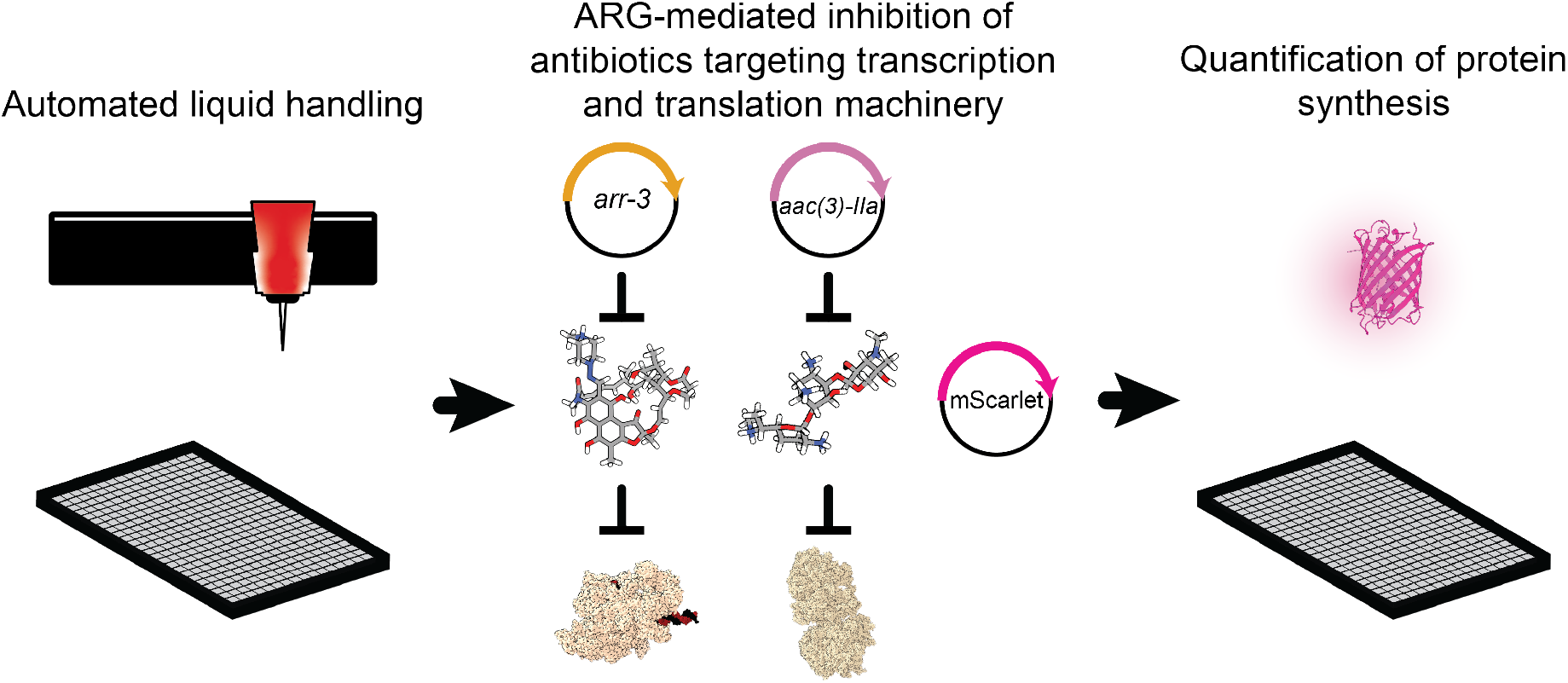

## INTRODUCTION

Antimicrobial resistance (AMR) is a global concern to human health, contributing to millions of deaths each year.^1^ Bacteria can acquire novel AMR through intrinsic genetic changes, including chromosomal mutation and recombination, as well as horizontal gene transfer (HGT), with the latter enabling rapid dissemination of antimicrobial resistance genes (ARGs) between strains and species.^2–4^ Genomic surveillance enables the identification of emerging ARGs and resistance-associated mutations; however, distinguishing between genes that confer clinically relevant resistance and sequence-similar homologues with negligible impact remains a bottleneck, often requiring labour-intensive genotype-to-phenotype characterisation.^3,5^ It can also be difficult to assess the relative contributions of different AMR mechanisms, for example, drug target mutations, efflux, and enzymatic modifications, which may all contribute to the overall resistance to a specific antibiotic or other therapeutic agent in bacterial cells.^6^

Here, we characterise two ARGs from distinct families as a proof-of-concept for profiling antibiotic-inactivating enzymes in *Escherichia coli* cell-free expression (CFE) systems. Enzymatic modification is one of the most important forms of resistance to aminoglycosides and is poorly understood given the large, diverse family of aminoglycoside-modifying enzymes able to act on these substrates.^2,10^ The gene *aac(3)-IIa* provides an example of this critical mechanism, encoding an aminoglycoside *N*-acetyltransferase that deactivates gentamicin, which otherwise inhibits protein synthesis by binding with the 30S ribosomal subunit.^11–13^ While a less common mechanism for rifampicin resistance, *arr-3* encodes an enzyme that catalyses ADP-ribosylation of rifampicin, impairing its ability to inhibit transcription via interaction with the RNA polymerase (RNAP) β-subunit.^8^ Both *aac(3)-IIa* and a close homologue of *arr-3* have been demonstrated to be active in cell-free lysates,^14–16^ and previous work has demonstrated the potential utility of CFE for screening ribosome-targeting inhibitors.^17^

Integrating CFE systems with automated workflows and biofoundries has enabled increased experimental throughput and reproducibility, with recent applications in biological engineering and natural product discovery; however, accessibility to biofoundries and large-scale automation remains limiting.^18,19^ In this study, we design a semi-automated workflow to characterise ARG activity in *E. coli* cell-free lysates with a compact benchtop Bio Molecular Systems Myra liquid handling robot that offers a low-cost alternative to large-scale automated systems. The ARGs *aac(3)-IIa* and *arr-3* are well-suited for developing this CFE-based methodology due to their direct, intracellular enzymatic activity on antibiotics and the activity of their cognate antibiotics being directed toward native transcription and translation machinery present in lysate-based CFE systems.^18^ This study adds to recent work developing automated and semi-automated CFE workflows using Opentrons OT-2 robotics^20^ and Echo Acoustic Liquid Handlers,^21^ with the focus of making CFE experimentation standardised, reproducible, and accessible to a broader range of users.

## RESULTS

### Automated liquid handling is equivalent to manual pipetting for CFE-based ARG activity assays

To establish the reliability of automated liquid handling for characterising ARG activity, we first compared results between automated and manual workflows. For initial characterisation, a plasmid harbouring *aac(3)-IIa* or a water control was added to a 384-well plate followed by the immediate addition of cell-free reaction mix, which included *E. coli* BL21 Star™ (DE3)pLysS lysate, supplemental substrates and co-factors known to maximise protein synthesis, and a plasmid encoding mScarlet, which was used as a quantitative readout of lysate protein synthesis (**Fig. 1a**). The cell-free reactions were incubated for 2 h to enable expression of *aac(3)-IIa* before the addition of gentamicin or water. Liquid transfers were designed for a total reaction volume of 15 μL, and the complete reactions were then incubated for 16 h. Expression of *aac(3)-IIa* protected cell-free reactions from gentamicin-dependent inhibition of protein synthesis, and no statistically significant differences (*n* = 9) were observed between manual and automated liquid dispensing across all conditions tested (**Fig. 1b**).

**Fig. 1:**
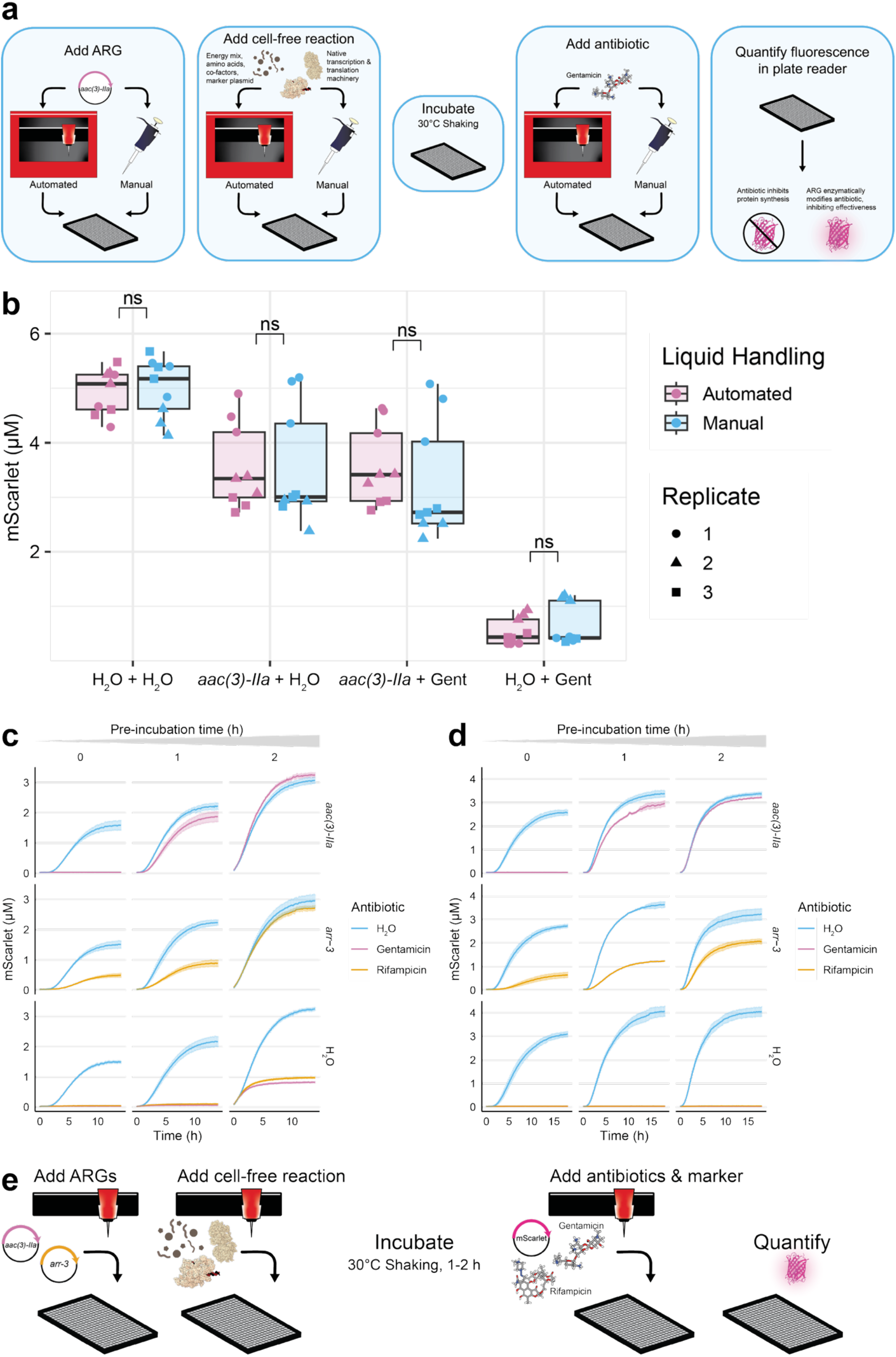
Automated liquid handling workflow for cell-free ARG expression functions equivalently to manual pipetting. **a)** Schematic for cell-free workflow comparing automated liquid handling to manual pipetting. Ribosome (PDB 4YBB), RNAP complex (PDB 6×43), and gentamicin (PubChem CID 3467) are shown for illustrative purposes, not to scale. **b)** mScarlet (μM) produced after 16 h across three independent replicates. Welch’s t-tests followed with Benjamini & Hochberg correction indicate no significant differences between manual and automated pipetting for each combination. **c, d)** ARG pre-incubation time determines protection efficacy in CFE antibiotic inhibition assays. mScarlet production over 16 h after ARG plasmids *aac(3)-IIa* and *arr-3* were either incubated with the cell-free reaction and mScarlet marker plasmid for 0-2 h prior to adding antibiotic (c) or incubated with the cell-free reaction but not the mScarlet marker plasmid for 0-2 h prior to adding both the antibiotic and mScarlet marker plasmid (d). **e)** Optimised automated liquid handling workflow. Plasmids, proteins, and antibiotics (rifampicin: PubChem CID 135398735) not shown to scale.

### Pre-incubation is essential for reproducible ARG activity in CFE assays

We rationalised that ARG activity within cell-free lysates would depend on a pre-incubation phase, allowing ARG synthesis prior to antibiotic-mediated inhibition of the RNAP or ribosomes by rifampicin and gentamicin, respectively. We tested different ARG pre-incubation durations in which mScarlet was either co-expressed during pre-incubation (**Fig. 1c**) or added after pre-incubation (**Fig. 1d**). We found that dependency on pre-incubation time differed between ARGs, with *aac(3)-IIa* requiring 1-2 h for optimal activity, while *arr-3* was at least partially active immediately. Co-expression of mScarlet during pre-incubation beyond 1 h resulted in detectable fluorescence under antibiotic treatments in the absence of ARGs, attributable to maturation of a small mScarlet pool synthesised before antibiotic-mediated inhibition (**Fig. 1c**). Adding the mScarlet plasmid after pre-incubation eliminated this background signal (**Fig. 1d**).

### Automated CFE quantifies aminoglycoside and rifampicin resistance gene activity

ARG-expressing cell-free reactions were challenged with increasing antibiotic concentrations to determine if the optimised semi-automated protocol could be used for robust quantification of enzymatic activity (**Fig. 1e**). Expression of *aac(3)-IIa* and *arr-3* notably increased the half-maximal inhibitory concentration (IC_50_) of gentamicin and rifampicin, respectively (**Fig. 2a**,**c**). Across three independent replicates, expression of *aac(3)-IIa* conferred a gentamicin IC_50_ value of 10.04 ± 0.84 µM, which was over 150-fold greater and statistically distinct (adjusted p-values < 5×10^-5^) from negative controls expressing either no ARG (55 ± 3.7 nM) or *arr-3* (64 ± 05.4 nM; **Fig. 2b**). Likewise, expression of *arr-3* conferred a rifampicin IC_50_ value (1.00 ± 0.13 µM) that was approximately 20-fold greater and statistically distinct (adjusted p-values < 5×10^-4^) from empty (63 ± 10 nM) and *aac(3)-IIa*-expressing controls (49 ± 14 nM; **Fig. 2d**). As predicted, IC_50_ values when *aac(3)-IIa* expression was challenged with rifampicin and *arr-3* was challenged with gentamicin did not statistically differ from null controls. Therefore, enzymes encoded by *aac(3)-IIa* and *arr-3* behave as expected in cell-free reactions, enzymatically modifying and inhibiting gentamicin or rifampicin,^14–16^ and provide a proof-of-concept for semi-automated characterisation of intracellular AMR activity in cell-free (**Fig. 2e**).

**Fig. 2:**
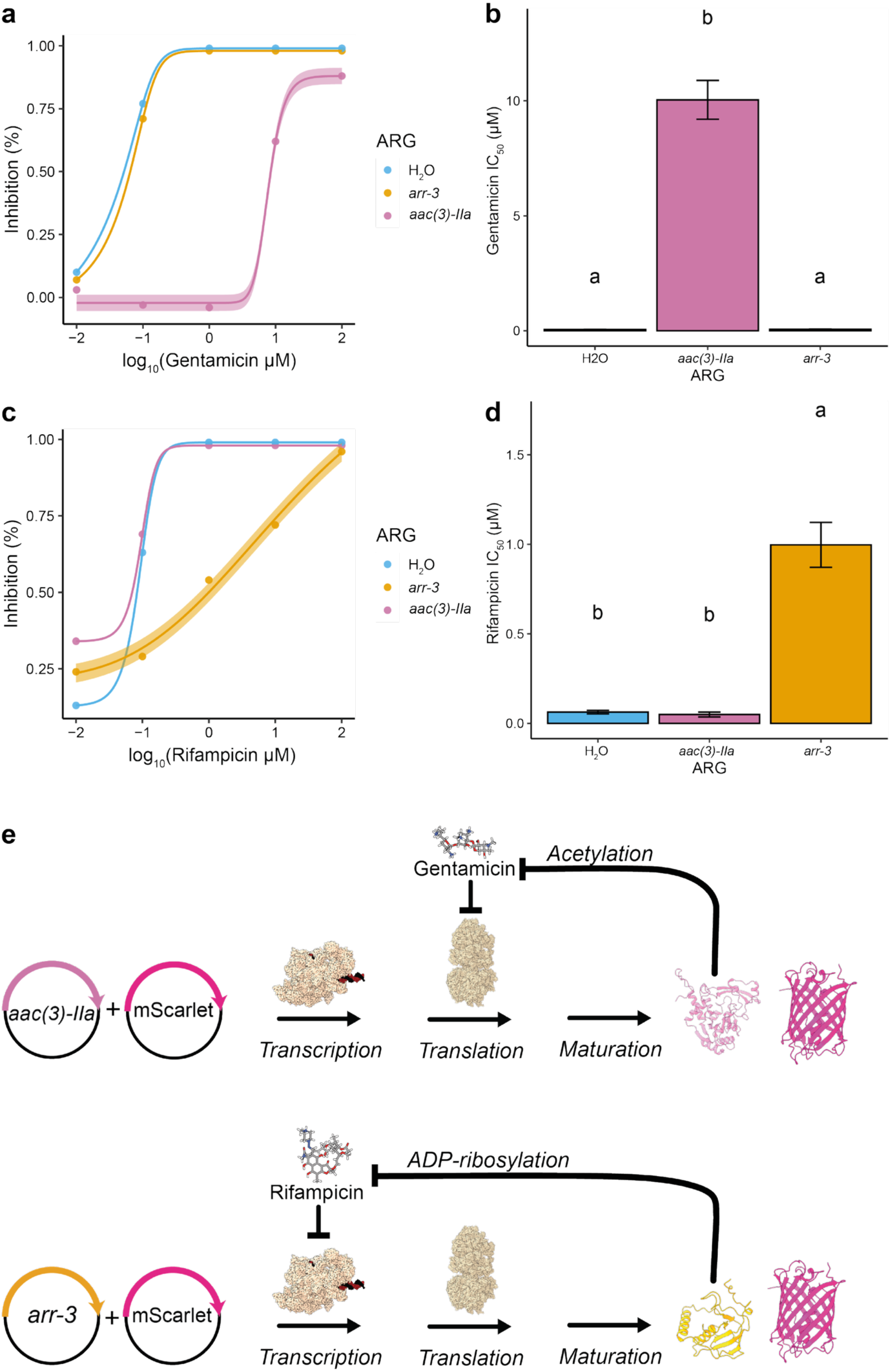
Cell-free quantification of *aac(3)-IIa* and *arr-3* activity. Cell-free expression of *aac(3)-IIa* and *arr-3* confers resistance to gentamicin and rifampicin, respectively. **a**,**c)** Percent inhibition of mScarlet cell-free protein synthesis for gentamicin (a) and rifampicin (c) treatments with points representing the median values of three technical replicates. **b)** A one-way ANOVA followed by TukeyHSD test indicates *aac(3)-IIa* mediates a statistically higher gentamicin IC_50_ value compared to *arr-3* and H_2_O controls. Mean values and standard error are shown for three independent replicates. **d)** A one-way ANOVA followed by TukeyHSD test indicates *arr-3* mediates a statistically higher rifampicin IC_50_ value compared to *aac(3)-IIa* and H2O controls. Mean values and standard error are shown for three independent replicates. **e)** Schematic showing hypothesized inhibitory mechanisms in cell-free. Drawings not to scale.

## DISCUSSION

In this study, we established a semi-automated CFE workflow with the Myra liquid handling system for the characterisation of ARGs. We validated the use of this automated system compared to skilled manual pipetting and optimised a 1-2 h pre-incubation period for robust ARG expression prior to antibiotic treatment. The requirement for a pre-incubation phase is supported by previous studies that indicate proteins in CFE systems undergo an initial phase of transcription that lasts up to 1 h for eGFP, a quasilinear protein synthesis phase for up to 8 h for eGFP, and a plateau upon completion.^22^ Maturation times are protein-dependent, causing these times to vary.^22,23^

While we cannot exclude the possibility that the disparity in pre-incubation times for *aac(3)-IIa* and *arr-3* is attributed to protein maturation kinetics, differing expression levels, or enzyme activity, we predict it reflects the distinct targets of the respective antibiotics within the gene expression pathway. Rifampicin targets RNAP-mediated transcription but does not prevent the translation of *arr-3* mRNA transcripts synthesized during a brief pre-incubation. Conversely, gentamicin inhibits ribosomal translation, meaning that *aac(3)-IIa* mRNA transcripts must be translated prior to antibiotic addition, necessitating a longer pre-incubation period. Additionally, the activity of a close homolog of Arr-3, Arr-2, has been shown to require NAD^+^.^14^ While all cell-free reactions were supplemented with NAD^+^, we did not investigate the relationship between NAD^+^ concentration and enzyme activity. For our purpose of developing a semi-automated workflow, a 1-2 h pre-incubation time sufficiently enabled detection of activity; however, future work characterising protein expression and enzymatic activity could further elucidate the discrepancy observed.

Scripts utilised for liquid handling are open-access and enable 384 unique 15-μL cell-free reactions on a single plate. Liquid transfer functioned with Myra default settings without concerns of bubbles causing variation in dispensed amounts before optimisation, as previously reported for various workflows.^20,21^ However, while Ekas *et al*. used an Echo Acoustic Liquid Handler, which is capable of transferring nL volumes, to screen cell-free reactions with a total volume of 1 μL, a limitation of the Myra and OT-2 robots is the minimum dispensing volume of 1 μL.^21^

Future work can utilise this accessible, semi-automated workflow to perform high-throughput characterisation of ARGs. Recent advancements with one-pot mutagenesis and linear DNA template expression in CFE systems could be employed for rapid phenotypic quantification of novel ARG activity with critical applications in healthcare.^24–27^ This addresses an important need to understand the contribution of specific ARGs to resistance for frontline therapeutics. While cell-free ARG characterisation is constrained to intracellular mechanisms, this limitation enables the determination of activity and substrate specificity for each ARG, presenting an alternative to whole-cell assays in which resistance could be achieved through any combination of efflux pump expression, import mechanisms, target modifications, or multiple ARGs.^6^ Understanding substrate specificity of ARGs is also important for new antibiotic discovery, especially where drugs are being designed and modified to evade known enzymatic modification, for example in resistance-breaker approaches, or for inhibitors targeting ARGs.^28–31^ Therefore, our cell-free approach for characterising ARGs may provide important applications in AMR surveillance and drug development.

## MATERIALS & METHODS

### Cell-free lysate preparation

Cell-free extracts of *E. coli* BL21 Star™ (DE3)pLysS were prepared according to a previously published workflow.^32^ Briefly, cells were streaked and grown on an LB agar plate overnight. A loopful of colonies was collected with a sterile loop and used to inoculate a 1-L culture of CFAI media (86 mM NaCl, 44 mM KH_2_PO4, 80 mM K_2_HPO4, 2.8 mM D-glucose, 11 mM D-lactose, 2 % (w/v) tryptone, 0.5 % (w/v) yeast extract, 0.6 % (v/v) glycerol, pH adjusted to 7.2 with KOH) in a 2-L glass baffled flask. The culture was incubated overnight at 30°C while shaking at 200 rpm. Cells were cooled on ice, pelleted (5000×*g*, 10 min, 4 °C), and resuspended in S30 buffer (10 mM Tris acetate pH 8.2, 14 mM Magnesium acetate tetrahydrate, 60 mM Potassium acetate, 2 mM dithiothreitol). Cells were pelleted again, and the supernatant was removed before resuspending the pellet in S30 buffer at a 1:1 (w/v) ratio. Extracts were lysed under the following sonication settings: 10 s pulse on, 10 s pulse off at 55-65% amplitude for an approximate active sonication time of 4 min. Lysed cells were centrifuged (15,000×*g*, 10 min, 4 °C), and the supernatant was collected, aliquoted and flash frozen in dry ice. A Bradford assay was used to estimate total protein concentration with Pierce™ Bradford Protein Assay Kit (Thermo Scientific) using bovine serum albumin as a standard, and extracts were diluted in S30 buffer to approximately 25-35 mg/mL for CFE.

### CFE

*E. coli* lysate-based cell-free reactions were comprised of the following: 248 μM CoA, 320 μM NAD, 731 μM cAMP, 65 μM folinic acid, 926 μM spermidine, 30 mM 3-PGA, 1.36 mM ATP, 1.36 mM GTP, 0.68 mM CTP, 0.68 mM UTP, 955 μM amino acids (except for leucine at 796 μM), 6 mM Mg-glutamate, 175 mM K-glutamate, 15 nM SP44-mScarlet-I plasmid (Addgene #163756), 3 % (w/v) PEG-8000, 48 mM Hepes buffer (pH 8.0), and approximately 10 mg/mL lysate.^33^ Plasmids encoding ARGs were added at final concentrations of 10-15 nM, and antibiotics were added at the specified concentrations after the specified pre-incubation period. A FLUOstar Omega (BMG Labtech) plate reader was used to measure relative mScarlet fluorescence over 16 h at 30°C, with double orbital shaking at 200 rpm between readings, and 584 nm/620-10 nm excitation/emission settings.

### Automated liquid handling

Unless otherwise noted, pipetting was performed with a Myra robot (Bio Molecular Systems) using the scripting interface. For 15 μL reactions, ARG plasmids were first added (120-180 nM 1.25 μL), followed by the cell-free reaction mixture with or without the mScarlet marker plasmid as specified (10-11.25 μL). After incubation at 30°C with shaking at 200 rpm for 0-120 min, antibiotics, which were pre-mixed with mScarlet at a 1:1 ratio if specified, were added to each well (1.25-2.5 μL). R scripts for generating the indexed sample input file and Myra-readable scripts to assign specified ARGs and antibiotics to each well are available on Github (https://github.com/Mo42-mb/Myra_automation.git).

### mScarlet standardisation

Purified mScarlet standards were used to convert relative fluorescence to μM concentration. For purification, *E. coli* BL21 Star™ (DE3)pLysS carrying pET-15b-6xHis-mScarlet-I (Addgene # 255536) was grown in 500 mL LB with 100 µg/mL carbenicillin at 37 °C, 200 rpm to an OD_600_ of 0.6, induced with 1 mM IPTG, and incubated at 16 °C, 200 rpm for 16–18 h. Cells were harvested (5000×*g*, 5 min, 4 °C) and resuspended in binding buffer (50 mM Tris-HCl pH 8.0, 500 mM NaCl, 10 mM imidazole, EDTA-free Pierce™ Protease Inhibitor (Thermo Scientific) per manufacturer’s instructions) in a 1:10 (w/v) ratio. Cells were lysed by sonication and clarified (15,000×*g*, 10 min, 4 °C). The clarified lysate was purified by gravity-flow Ni^2+^-Immobilized Metal Affinity Chromatography (IMAC) by loading on a 20-mL equilibrated column with Ni^2+^-charged Chelating Sepharose™ Fast Flow resin (Cytiva). Buffers with consistent Tris-HCl and NaCl compositions but increasing imidazole concentrations were used to wash (30 mM then 70 mM imidazole) and elute (400 mM imidazole) proteins. mScarlet was quantified by absorbance at 280 nm using an extinction coefficient predicted by ProtParam (ExPASy). RFU values were recorded for a standard series (0-8 μM) with three technical replicates and three independent replicates, and a linear regression (Adjusted R^2^ = 0.9999) was used to convert all experimental RFU values to μM concentration (**Supplementary Fig. S1**).

### Statistical analyses

All analysis was performed in R. For each independent replicate, the median of three technical replicates was calculated to reduce the effect of outliers. Data was evaluated for outliers, normality, and homogeneity of variances prior to performing Welch’s t-tests with Benjamini & Hochberg correction (rstatix; 10.32614/CRAN.package.rstatix) or a one-sided ANOVA followed by a TukeyHSD test (α = 0.05).

### IC_50_ characterisation

Percent inhibition was calculated as the difference between the mScarlet concentration for the dH_2_O control and the mScarlet concentration under the specified antibiotic treatment, divided by the mScarlet concentration of the dH_2_O control. N-parameter regressions were calculated and visualised with the nplr package in R, with the goodness of fit for each prediction being at least 0.992 (10.32614/CRAN.package.nplr).

### Plasmids

ARG plasmids were developed with EcoFlex Golden Gate Assembly using the J23100 promoter, pET RBS, BB0012 terminator, and pTU1-B backbone.^34^ Coding sequences of *aac(3)-IIa* (Addgene #255535) and *arr-3* (Addgene #255534) expression vectors were derived from NCBI X13543 and NCBI AJ277027, respectively, and *Klebsiella pneumoniae* codon-optimised. All plasmids used for CFE were purified using a QIAGEN Plasmid Midi Kit. DNA was resuspended in 1:1 (v/v) nuclease-free H_2_O: isopropanol with 149 mM sodium acetate and incubated at −20 °C overnight. DNA was pelleted (10,000×*g*, 2 min), washed three times with 70% ethanol, vacuum-dried (∼10 min, 60 °C), and resuspended in nuclease-free H_2_O. Concentration was measured using a Qubit Fluorometer.

## ASSOCIATED CONTENT

### Material Availability Statement

All plasmids used in this work have been deposited on Addgene (#163756, #255534-6).

### Data Availability Statement

All data generated or analysed during this study is provided on Figshare (doi:10.6084/m9.figshare.32055030).

### Supporting Information

Quantification of mScarlet with purified standards (Supplementary Fig. S1).

### Author Contributions

S.J.M., M.B., and J.M.S. conceived and designed the research. M.B. developed methodology, conducted experiments, and performed analysis/visualisation. M.B. wrote the original draft, and S.J.M., J.M.S., and B.M. provided editing. S.J.M and M.S. acquired funding and provided resources.

### Funding

M.B. and B.M. were supported by the BBSRC awarded to S.J.M (BB/Y005074/2) and J.M.S (BB/Y005325/1).

## Acknowledgements

Thank you to Alexandra-Georgiana Butulan and Charlotte A. Woolley for their valuable feedback throughout this project. We would also like to thank Mike Slinger from Bio Molecular Systems for technical support for the Myra liquid handling robot.

## CITATIONS

1. Naghavi, M. et al. Global burden of bacterial antimicrobial resistance 1990–2021: a systematic analysis with forecasts to 2050. The Lancet 404, 1199–1226 (2024).

2. Christaki, E., Marcou, M. & Tofarides, A. Antimicrobial Resistance in Bacteria: Mechanisms, Evolution, and Persistence. J. Mol. Evol. 88, 26–40 (2020).

3. Djordjevic, S. P. et al. Genomic surveillance for antimicrobial resistance — a One Health perspective. Nat. Rev. Genet. 25, 142–157 (2024).

4. Urban-Chmiel, R. et al. Antibiotic Resistance in Bacteria—A Review. Antibiotics 11, 1079 (2022).

5. Zhang, A.-N. et al. An omics-based framework for assessing the health risk of antimicrobial resistance genes. Nat. Commun. 12, 4765 (2021).

6. Darby, E. M. et al. Molecular mechanisms of antibiotic resistance revisited. Nat. Rev. Microbiol. 21, 280–295 (2023).

7. Elias, R. et al. Dissemination of arr-2 and arr-3 is associated with class 1 integrons in Klebsiella pneumoniae clinical isolates from Portugal. Med. Microbiol. Immunol. (Berl.) 214, 6 (2025).

8. Surette, M. D., Spanogiannopoulos, P. & Wright, G. D. The Enzymes of the Rifamycin Antibiotic Resistome. Acc. Chem. Res. 54, 2065–2075 (2021).

9. Pradier, L. & Bedhomme, S. Ecology, more than antibiotics consumption, is the major predictor for the global distribution of aminoglycoside-modifying enzymes. eLife 12, e77015 (2023).

10. Zárate, S. et al. Overcoming Aminoglycoside Enzymatic Resistance: Design of Novel Antibiotics and Inhibitors. Molecules 23, 284 (2018).

11. Allmansberger, R., Briu, B. & Piepersberg, W. Genes for gentamicin-(3)-N-acetyl-transferases III and IV. Mol. Gen. Genet. MGG 198, (1985).

12. Ramirez, M. S. & Tolmasky, M. E. Aminoglycoside modifying enzymes. Drug Resist. Updat. 13, 151–171 (2010).

13. Hutchings, M. I., Truman, A. W. & Wilkinson, B. Antibiotics: past, present and future. Curr. Opin. Microbiol. 51, 72–80 (2019).

14. Quan, S. et al. ADP-Ribosylation as an Intermediate Step in Inactivation of Rifampin by a Mycobacterial Gene. Antimicrob. Agents Chemother. 43, 181–184 (1999).

15. Morisaki, N. et al. Structures of ADP-Ribosylated Rifampicin and Its Metabolite: Intermediates of Rifampicin-ribosylation by Mycobacterium smegmatis DSM43756. J. Antibiot. (Tokyo) 53, 269–275 (2000).

16. Maurer, F. P. et al. Aminoglycoside-modifying enzymes determine the innate susceptibility to aminoglycoside antibiotics in rapidly growing mycobacteria. J. Antimicrob. Chemother. 70, 1412–1419 (2015).

17. Höger, B., Peifer, C. & Beitz, E. Cell-free production of fluorescent proteins for the discovery of novel ribosome-targeting antibiotics. J. Microbiol. Methods 213, 106814 (2023).

18. Rice, A. J. et al. Cell-free synthetic biology for natural product biosynthesis and discovery. Chem. Soc. Rev. 54, 4314–4352 (2025).

19. Jun, J.-S., Hong, S., Park, J.-H., Shin, J. & Lee, D.-H. Automated and Programmable Cell-Free Systems for Scalable Synthetic Biology with a Focus on Biofoundry Integration. J. Microbiol. Biotechnol. 35, e2507019 (2025).

20. Brown, D. M. et al. Semi-automated Production of Cell-free Biosensors. bioRxiv (2024).

21. Ekas, H. M. et al. An Automated Cell-Free Workflow for Transcription Factor Engineering. ACS Synth. Biol. 13, 3389–3399 (2024).

22. Garenne, D. et al. Cell-free gene expression. Nat. Rev. Methods Primer 1, 49 (2021).

23. Balleza, E., Kim, J. M. & Cluzel, P. Systematic characterization of maturation time of fluorescent proteins in living cells. Nat. Methods 15, 47–51 (2018).

24. Norouzi, M., Zayeni, R., Singh, S. & Pardee, K. Cell-Free Dot Blot as a Practical and Adaptable Immunoassay Platform for the Detection of Antibody Response in Human and Animal Sera. J. Vis. Exp. 67973 (2025) doi:10.3791/67973.

25. Sato, W. et al. One-pot cloning and protein expression platform for genetic engineering. bioRxiv https://www.biorxiv.org/content/10.1101/2025.08.28672974v1. (2025).

26. Pandi, A. et al. Cell-free biosynthesis combined with deep learning accelerates de novo-development of antimicrobial peptides. Nat. Commun. 14, 7197 (2023).

27. Yim, S. S., Johns, N. I., Noireaux, V. & Wang, H. H. Protecting Linear DNA Templates in Cell-Free Expression Systems from Diverse Bacteria. ACS Synth. Biol. 9, 2851–2855 (2020).

28. Laws, M., Shaaban, A. & Rahman, K. M. Antibiotic resistance breakers: current approaches and future directions. FEMS Microbiol. Rev. 43, 490–516 (2019).

29. Alanzi, A. R., Alqahtani, M. J., Alqahtani, J. H. & Alharbi, H. A. A computational study exploring echinoderm-derived compounds for inhibition of aminoglycoside acetyltransferases. PLOS One 20, e0327409 (2025).

30. Chiem, K. et al. Inhibition of Aminoglycoside 6′-N -Acetyltransferase Type Ib-Mediated Amikacin Resistance in Klebsiella pneumoniae by Zinc and Copper Pyrithione. Antimicrob. Agents Chemother. 59, 5851–5853 (2015).

31. Boehr, D. D. et al. Broad-Spectrum Peptide Inhibitors of Aminoglycoside Antibiotic Resistance Enzymes. Chem. Biol. 10, 189–196 (2003).

32. Smith, P. E. J., Slouka, T., Dabbas, M. & Oza, J. P. From Cells to Cell-Free Protein Synthesis within 24 Hours Using Cell-Free Autoinduction Workflow. J. Vis. Exp. 62866 (2021) doi:10.3791/62866.

33. Sun, Z. Z. et al. Protocols for Implementing an Escherichia coli Based TX-TL Cell-Free Expression System for Synthetic Biology. J. Vis. Exp. 50762 (2013) doi:10.3791/50762.

34. Moore, S. J. et al. EcoFlex: A Multifunctional MoClo Kit for E. coli Synthetic Biology. ACS Synth. Biol. 5, 1059–1069 (2016).

