## Supplementary Fig. S1 for "A liquid handling platform for standardised quantification of cell-free enzymatic activity encoded by antimicrobial resistance genes"

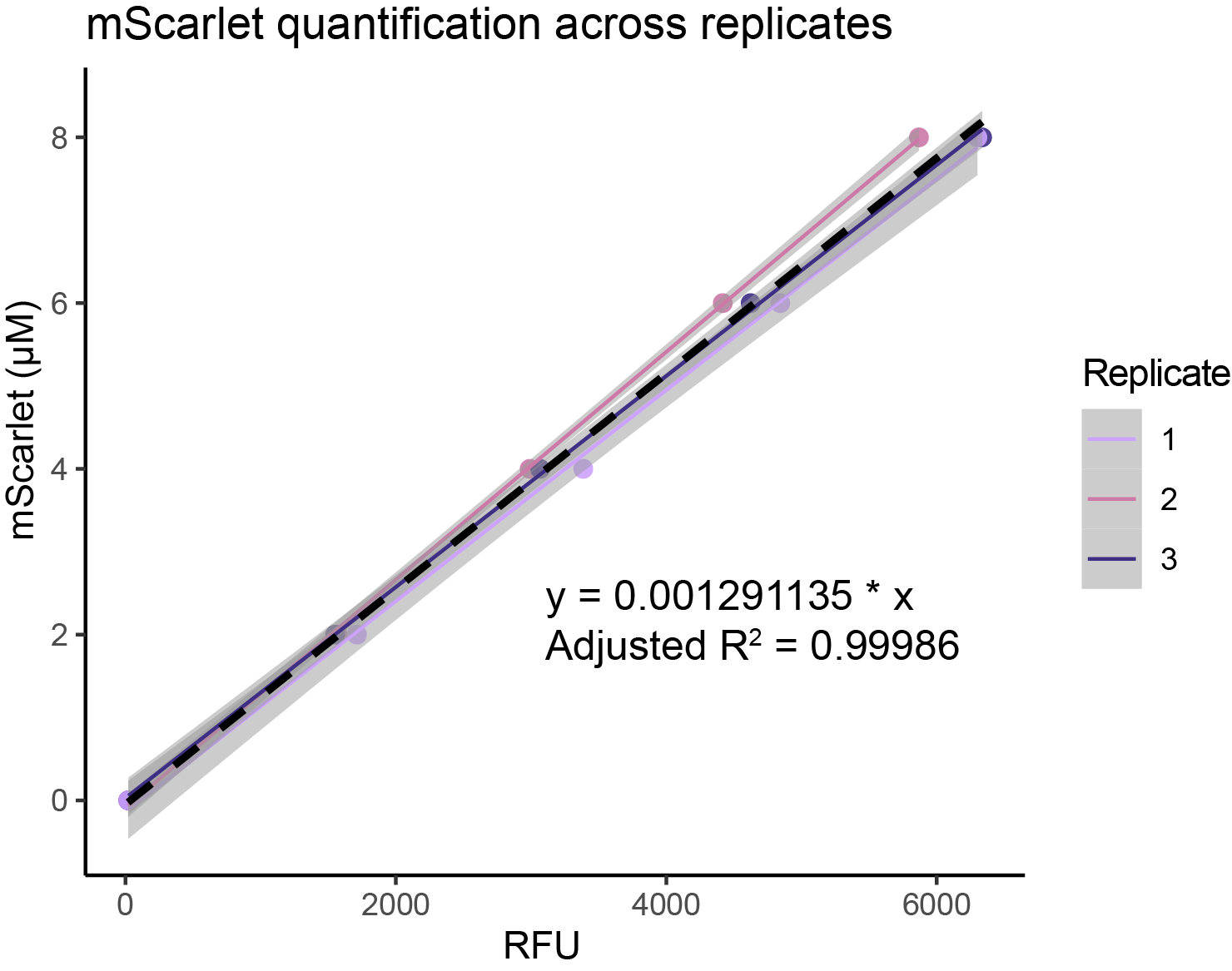

**Supplementary Fig. S1: Linear relationship between relative fluorescence units (RFU) and mScarlet concentration for five known standards across three independent replicates.** The average linear regression is shown in a black dashed line with a grey ribbon representing standard error.
